# Hydrocarbon-degrading microbial populations in permanently cold deep-sea sediments in the NW Atlantic

**DOI:** 10.1101/2024.05.16.593224

**Authors:** Oyeboade Adebayo, Srijak Bhatnagar, Jamie Webb, Calvin Campbell, Martin Fowler, Natasha M. MacAdam, Adam Macdonald, Carmen Li, Casey R.J. Hubert

**Affiliations:** Department of Biological Sciences, University of Calgary, AB T2N 1N4, Canada; Applied Petroleum Technology (Canada), Calgary, AB T2N 1Z6, Canada; Geological Survey of Canada-Atlantic, Dartmouth, NS B3B 1A6, Canada; Nova Scotia Department of Natural Resources and Renewables, Halifax, NS B2H 4G8, Canada; Faculty of Science and Technology, Athabasca University, Athabasca, AB T9S 3A3, Canada

**Keywords:** deep sea sediments, hydrocarbon biodegradation, microbial community composition, Gammaproteobacteria

## Abstract

Permanently cold deep-sea sediments (2500-3500 m water depth) with or without indications of thermogenic hydrocarbon seepage were exposed to naphtha to examine the presence and potential of aerobic hydrocarbon-degrading microbial populations. Monitoring these microcosms for volatile hydrocarbons by GC-MS revealed sediments without *in situ* hydrocarbons responded more rapidly to naphtha amendment than hydrocarbon seep sediments overall, but seep sediments removed BTEX compounds more readily. Naphtha-driven aerobic respiration was more evident in surface sediment (0-20 cmbsf) than deeper anoxic layers (>130 cmbsf) that responded less rapidly. In all cases, enrichment of Gammaproteobacteria included lineages of *Oleispira*, *Pseudomonas*, and *Alteromonas* known to be associated with marine oil spills. On the other hand, taxa known to be prevalent *in situ* and diagnostic for thermogenic hydrocarbon seepage in deep sea sediment did not respond to naphtha amendment. This suggests a limited role for seep-associated populations in the context of oil spill biodegradation.

## 1. Introduction

Offshore oil exploration has been happening for over 100 years (Hyne, 2001) with recent advances in drilling technology seeing activities extending farther offshore into deeper waters (EIA, 2016). Ultra deep-water operations in the Gulf of Mexico include Perdido at 2,450 m and Stones at 2,900 m, with similarly deep discoveries off the coast of Brazil such as the Carcana site in 2,030 m (Offshore Technology, 2017). Deep water presents challenging operational environments as illustrated by the Deepwater Horizon (DWH) oil blowout that occurred while producing oil in approximately 1,500 m water (Hazen et al., 2010; Camilli et al., 2011; Shukla and Karki 2016). This highlights the importance of understanding the microbial ecology of the deep sea, both with respect to baseline microbial communities (Joye, 2015; Ferguson et al., 2023) and the potential these microbiomes harbour for the biodegradation of spilled oil.

Hydrocarbonoclastic bacteria in marine ecosystems can derive carbon and energy from the degradation of petroleum hydrocarbons (Hazen et al., 2010, Kimes et al, 2014, Yang et al., 2016, Berry and Gutierrez, 2017). These bacteria have been observed to proliferate following oil spills and thus represent catalytic potential that can be harnessed for bioremediation (Yakimov et al., 2007, Acosta-Gonzalez et al., 2015, Joye et al., 2016, Duran and Cravo-Laureau, 2016; Yang et al., 2016). Reasons for the presence of oil-degrading microbial populations in the ocean include widespread occurrences of natural seabed hydrocarbon seepage (Head et al., 2006), yet it is unclear whether bacteria commonly understood to be hydrocarbonoclastic (Berry and Gutierrez, 2017; Gutierrez et al., 2013; Sanni et al., 2015) are a guild that overlaps with dominant microbial populations inhabiting hydrocarbon seep sediments in the deep sea (Dong et al., 2019, 2020; Chakraborty et al., 2020; Li et al., 2023).

Most published studies following the DWH interrogated the response of microbial communities in the water column to the introduction of spilled oil and gas (Hazen et al., 2010; Redmond and Valentine., 2012; Yang et al., 2016). Additional research showed 2-15% of the oil released from the Macondo wellhead eventually became deposited onto the deep-sea sediments via marine oil snow sedimentation and floc accumulation (Passow et al., 2012; Valentine et al. 2014; Chanton et al. 2015). Hydrocarbons deposited in marine sediments become absorbed into the sediment organics impacting ecosystem functioning (Karickhoff et al., 1979; Eadie et al., 1982, McGroddy and Farrington., 1995; Coates et al., 1997, Cravo-Laureau and Duran., 2014). Nearby sediments investigated following the DWH blowout revealed oil deposition penetrating the top 5 cm of the seabed (Joye et al., 2014). Natural hydrocarbon seeps on the other hand receive inputs of petroleum compounds from below via slower advection as part of geological petroleum systems (Joye, 2020). As a result, deep sea sediments experiencing hydrocarbon seepage are enriched in particular taxa, including *Caldatribacteriota*, *Aminicenantes* and *Campilobacterota* (Chakraborty et al. 2020; Li et al. 2023). Whether or not these taxa would be involved in the biodegradation of petroleum compounds entering the seabed from above in an oil spill scenario requires further investigation.

Temperature is another factor that controls the fate of oil in the marine environment. Physical and chemical properties of hydrocarbon compounds as well as the metabolic rates catalyzed by hydrocarbonoclastic microbes are influenced by temperature. The deep sea is generally very cold, with temperatures close to 0℃ (Yasuhara and Danovaro 2016). To investigate the potential for psychrophilic aerobic hydrocarbon biodegradation in a deep-sea setting, sediments with and without in situ hydrocarbons were incubated with naphtha (low molecular weight hydrocarbons including short- and long-chain alkanes and monoaromatic hydrocarbons) at 4°C for 100 days and compared with each other to test the hypothesis that cold seep sediments are primed for biodegradation.

## 2. Materials and methods

### 2.1. Seismic description

Seismic interpretation was performed on the Shelburne (NS24-S006-003E) and Tangier (NS24-B071-001E) 3D surveys using Schlumberger’s Petrel software platform to assess potential piston coring locations based on amplitude anomalies and the presence of direct hydrocarbon indicators (DHIs). Geophysical Reports for the 3D seismic surveys can be accessed here: https://cnsopbdigitaldata.ca/dmc-summary/. DHIs were inferred to be associated with possible seabed seeps on the basis of potential hydrocarbon migration pathways in the form of faults from deeper in the subsurface up to the seafloor (Figure 1B, C).

**Figure 1:**
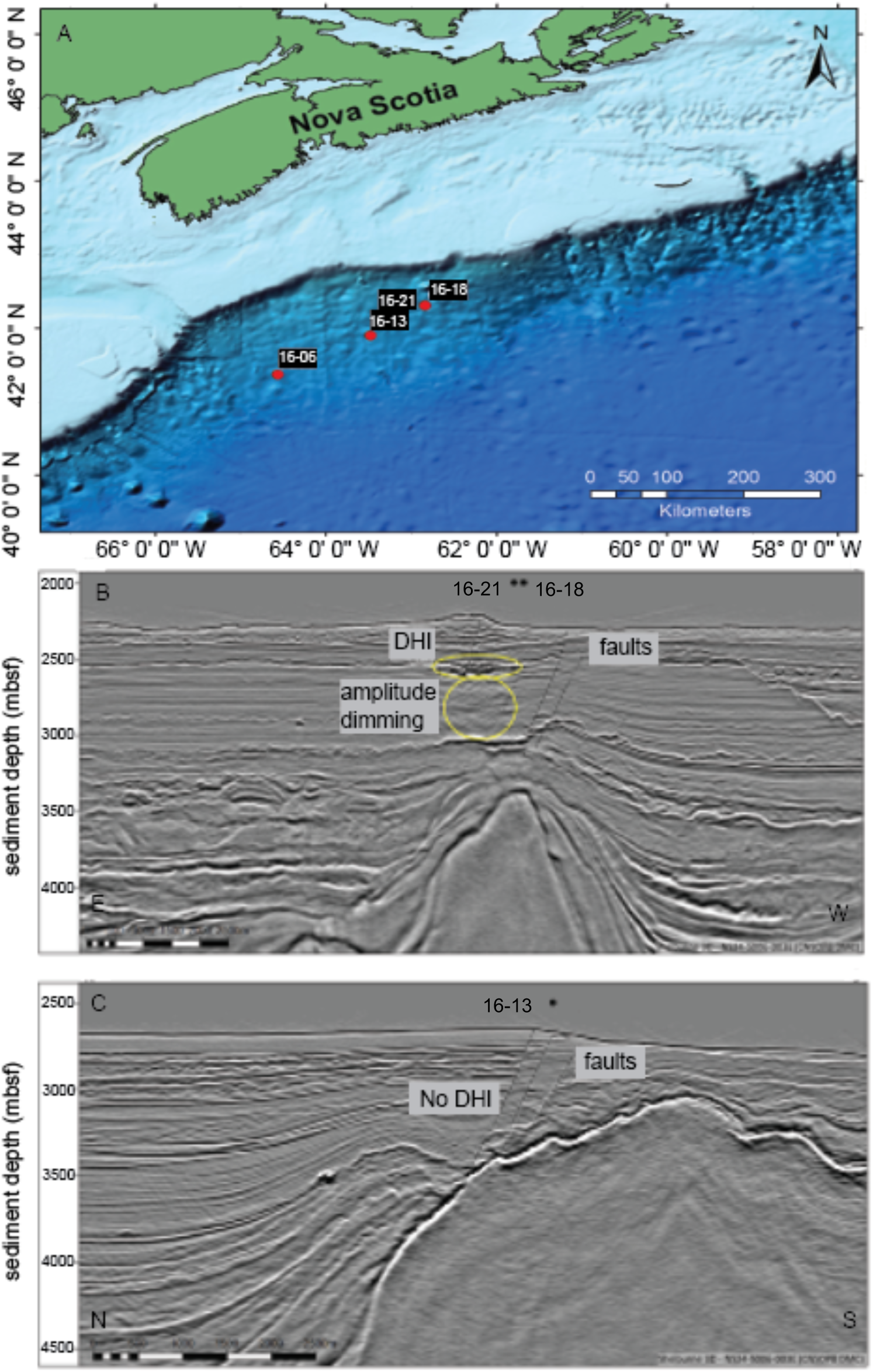
Map of the Scotian Slope, NW Atlantic Ocean showing (A) the location of four piston coring sites sampled aboard CCGS *Hudson* in 2016. Sites 16-18 and 16-21 are 160m apart and overlap on the map. Geophysical images showing sites 16-18 and 16-21 (B) and site 16-13 (C) at higher resolution reveal suspected direct hydrocarbon indicators (DHIs) such as deep-seated fault near surface, amplitude diming (in circles) and faults (broken lines) for sites 16-18 and 16-21, but not 16-13 which shows only faults to surface above a large shallow diapir.

### 2.2. Study site and sampling

Sampling was conducted onboard the CCGS *Hudson* in June and July in 2016. Sediments from four stations 16-06, 16-13, 16-18 and 16-21 were sampled via piston coring along the Scotian slope off the coast of Nova Scotia in the NW Atlantic Ocean (Fig 1). Within a few hours of core retrieval, sediments from these cores were sub-sampled and used to establish experimental microcosms that were amended with hydrocarbons and incubated at 4°C. Surface sediments (0-20 cmbsf) were sampled from 16-06, 16-13 and 16-18, and deeper sediments were also subsampled at 134-141 cmbsf (16-18) and 142-148 cmbsf (16-21).

### 2.3. Sulfate measurements

Sulfate concentrations in sediment porewater were measured at several depths following porewater extraction by centrifugation of a small aliquot of wet sediment (ca. 500 mg) taken from cores that were longitudinally sectioned on board the ship. Porewater was obtained by centrifugation. Initial sulfate measurements were made onboard by monitoring barium sulfate using the “USEPA SulfaVer 4” method with the Pocket Colorimeter II (Hach, Canada) and barium chloride ampules (AccuVac Ampules, Hach Canada). This rapid estimation of sulfate profiles and hence overall redox zonation in the sediments guided the choice of deeper sediment layers from cores 0018 and 0021. Additional sediment aliquots sampled in parallel were immediately stored on board in a -20°C freezer. These samples were eventually thawed and centrifuged similarly, allowing for sulfate measurements using a Dionex ICS 5000 reagent-free ion chromatography system (Thermo Scientific, CA, USA) equipped with an anion-exchange column (Dionex IonPac AS22; 4 × 250 mm; Thermo Scientific, USA), an EGC-500 eluent generator cartridge, and a conductivity detector. The eluent was Na_2_CO_3_ (4.5 mM) and NaHCO_3_ (1.4 mM) with a flow rate of 1.2 mL min^−1^ at 30 °C column temperature.

### 2.4. Hydrocarbon gas and liquid analysis in piston core sediment samples

Sediment was collected immediately after core retrieval from near the base of each core for hydrocarbon gas analysis by placing 200-300 g of sediment into a 500 ml Isojar that was flushed with nitrogen before sealing. Headspace gas aliquots were transferred to exetainers as 0.1-1.0 ml, arranged in a Gerstel MPS2 autosampler and injected into an Agilent 7890 RGA GC equipped with Molsieve and Poraplot Q columns, a flame ionisation detector (FID) and a thermal conductivity detector (TCD). Hydrocarbons were measured by FID. To measure liquid hydrocarbons, separate samples were wrapped in aluminium foil and stored at -20℃ on board. Following extraction of organic matter, liquid hydrocarbons from this fraction were measured using a HP7890 A GC instrument equipped with a CP-Sil-5 CB-MS column (length 30 m, i.d. 0.25 mm, film thickness 0.25 μm) using synthesized C_20_D_42_ compound as an internal standard.

### 2.5. Sediment microcosms amended with low-molecular weight hydrocarbons

Microcosms set up under oxic conditions (i.e., with air in the headspace) were established to assess how aerobic microbial communities in different surface sediments respond to exogenous hydrocarbon exposure. Sediment from 0-20 cmbsf was sampled from cores 16-06, 16-13 and 16-18. Deeper sediment layers corresponding to the sulfate reduction zone were sampled from cores 16-18 (134-141 cmbsf) and 16-21 (142-148 cmbsf), to additionally test for the presence of more deeply buried aerobic microbial communities and their ability to respond to hydrocarbon exposure in oxic microcosms. Microcosms consisted of 6 ml of sediment and 25 mL of ONR7a medium (Dyksterhouse et al., 1995) added to sterile 50 mL serum bottles with air in the headspace that were sealed with septa and aluminum crimp tops. Bottles were amended with 0.2% (v/v) naphtha, which is a qualitative reference blend of low molecular weight alkanes and aromatics. Heat killed controls (autoclaved microcosms), unamended controls (artificial seawater and sediment without naphtha amendment) and uninoculated controls (no sediment) were also established and incubated in parallel. All microcosms were kept static and incubated as triplicate bottles in the dark at 4°C for between 100 and 106 days.

### 2.6. Microbial respiration in microcosms

Headspace carbon dioxide and oxygen levels were measured at different sampling points using an Agilent 7890B gas chromatograph equipped with a thermal conductivity detector (TCD) and according to a protocol described elsewhere (Novotnik et al. 2019). The instrument operated under the following parameters: TCD temperature: 200°C, reference flow: He 40 mL/min. FID: Heater T: 200°C, Air flow 400 ml/min, H_2_ fuel flow 50 ml/min. Column 1: 0.5 m × 1/8″ Hayesep N (80/100 mesh); Column 2: 6′ × 1/8″ Hayesep N (80/100 mesh); Column 3: 8′ × 1/8″ MS5A (60/80 mesh). A reference gas mix from Praxair (Mississauga, Canada) was used for calibration.

### 2.7. Hydrocarbon consumption in microcosms

Volatile hydrocarbons were analyzed by injecting 100 µl headspace sample from each microcosm directly into an Agilent 6890N gas chromatograph/mass spectrometer (GC/MS) with a model 5973 inert mass selective detector and HP-5MS capillary column (30 m length, 0.25 mm internal diameter and 0.25 µm film thickness). The injector temperature was 250°C, and He carrier gas had a flow rate of 1 ml min^-1^ and split/splitless ratio of 1:10. The GC temperature program was 40°C for 5 min, then ramping up at 5°C min^-1^ to 85°C (Abu Laban et al., 2015). The GC/MS was run in full scan monitoring mode for *m*/*z* = 50-500. Headspace hydrocarbon depletion was assessed using the GC-MS method described by Prince and Suflita (2007) using 1,1,3-trimethylcyclohexane as the conserved internal standard (Townsend et al., 2004). The percentage of specific hydrocarbon (HC) compounds remaining in the headspace was calculated as (A_sample_/C_sample_)/(A_heat killed_/C_heat killed_) × 100, where A and C represent specific HC compounds and an internal standard respectively (Tan et al., 2015).

### 2.8. DNA extraction and 16S rRNA gene sequencing

Total genomic DNA was extracted from microcosm sediment slurries using the DNeasy PowerSoil kit (Qiagen, Valencia, CA, USA) according to manufacturer protocols. Extracted DNA was quantified using a Qubit 2.0 Fluorometer (Invitrogen). The V3–V4 hypervariable region of Bacterial and Archaeal 16S rRNA genes was amplified by polymerase chain reaction (PCR) using the universal primers Pro341F and Pro805R (Takahashi et al., 2014). Amplicon libraries were generated as triplicate twenty-five microliter reactions of 5-10 ng/μL DNA template, 1 μM of each primer, and 12.5 μL 2x KAPA HiFi Hot Start (Kapa Biosystems, Boston, USA). Amplification was performed using a Nexus GSX1 Master cycler (Eppendorf, Germany) as follows: initial denaturation at 94°C for 2 min, followed by 35 cycles of denaturation at 94°C for 30 sec, annealing at 58°C for 60 sec, and extension at 72°C for 60 sec, and final elongation at 72°C for 5 min. Triplicate PCR products were pooled and prepared for Illumina paired end sequencing using Illumina’s dual indexing protocol. Sequencing was performed using an in-house benchtop Illumina MiSeq sequencer (Illumina, San Diego, CA, USA).

### 2.9. DNA sequence analysis

Primer and adapter removal of raw demultiplexed reads was performed using cutadapt v1.16 (Martin, 2011). Primer trimmed sequences were quality checked and merged in DADA2 v1.9.0 (Callahan et al., 2016). Using the filterAndTrim function in DADA2, forward and reverse reads were trimmed to 280 bp and 220 bp, with filtering for reads with no ambiguous bases (maxN=0). Any reads with a quality score below 8 were truncated (trunqQ = 8). All suspect phiX sequences from the Illumina run were removed (rm.phix=TRUE) and only reads with expected error less than 2 for forward and 4 for reverse reads, were retained (max EE = c(2,4)). Two million high quality random forward and reverse reads that passed these filters were used to learn the error rates. Using error profiles of forward and reverse reads, libraries were merged using the mergePairs function in DADA2. From these merged reads amplicon sequence variants (ASVs) were inferred using makeSequenceTable function, followed by chimera removal using removeBimeraDenovo function. Chimera-free SVs were then assigned taxonomy using assignTaxonomy function with Silva nr training set v128 and a minimum bootstrap of 80. The deepest assigned taxonomy of each sequence variant was chosen to depict the taxonomic classification.

The ASV table generated was then consolidated at the genus level and used in Divnet (Willis and Martin, 2022) within the R software package to calculate alpha diversity indices with the parameters, seed=2021 and base=ASV 46. Differential abundance analysis was performed on a table with all ASVs using *differentialTest* function of Corncob (Martin et al 2020). With a seed set to 2021 reads, all differential abundance analyses were carried out using Wald test with bootstrapping and a false detection rate of 0.05, for each experimental parameter and combinations (e.g., hydrocarbon amendment, comparison between cores, etc). Using this approach, a set of statistically significant differentially abundant ASVs was determined. The abundance of these ASVs was plotted using package ggplot2 v 2.2.1 (Wickham 2016).

Sequences for six ASVs suspected to belong to genera considered to be obligate hydrocarbonoclastic bacteria (Yakimov et al., 2007) were used in BLAST searches of the NCBI nr database. Hits with ≥99% sequence identity to these six 16S rRNA gene sequences were compiled and aligned using SINA aligner, (Pruesse et al., 2012) with *Sulfolobus islandicus* (AY247900.1) as an outgroup. Phylogenetic tree reconstruction used RaxML (Stamatakis 2014) with Gamma livelihood and GTR models, and iTOL for visualization.

### 2.10. Data availability

Raw sequencing data used in this study can be accessed through NCBI BioProject Acc# PRJNA1014963

## 3. Results

### 3.1: Geophysical and geochemical evidence of hydrocarbon seepage

Geophysical imagery revealed sites 16-18 and 16-21 have subsurface acoustic features considered to be direct hydrocarbon indicators (DHIs), faulting, and a seafloor irregularity (Figure 1; Table 1). Site 16-13 exhibits faults to surface but no apparent DHI (Fig 1C). In agreement with these geophysical observations, the strongest geochemical evidence for sedimentary hydrocarbon gases and liquids was observed in cores 16-18 and 16-21 (Fig 2A, B). Furthermore, downcore sulfate concentration profiles differed distinctly between sampling sites (Fig 2C). A much steeper drop in sulfate concentration was observed in cores 16-18 and 16-21, where sulfate was completely depleted by 250 cmbsf; this is consistent with hydrocarbons providing additional substrates elevated sulfate reduction in these sediments. In contrast, cores 16-06 and 16-13 exhibited no marked drop in sulfate, with relatively flat profiles throughout the top 500 cm of the seabed.

**Figure 2:**
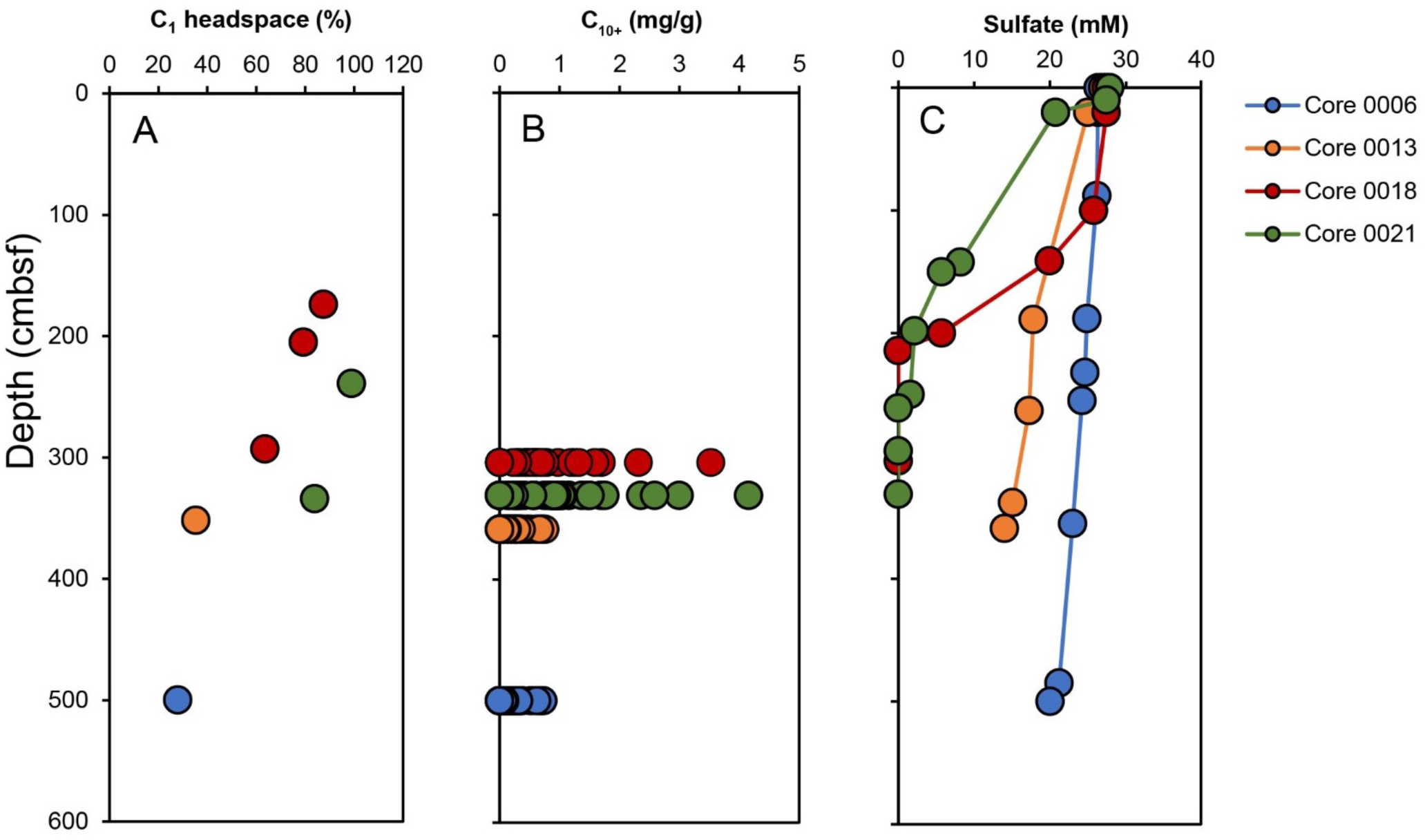
Sediment geochemical parameters in four sediment cores. (A) C_1_ gas measurements made following incubating different sediment depths in isojars immediately after sampling. (B) Liquid hydrocarbons were assessed using GC-MS for sediment sampled from the bottom of each of the cores, with each symbol of the same colour denoting different individual compounds in the C_10_ to C_42_ range. (C) Porewater sulfate concentrations determined at different depths throughout each core.

**Table 1:** Site location and core information of sampled cores^1^ from the Scotian slope, NW Atlantic Ocean.

| **Core** | **Coordinates** | **Water depth**<br>**(m)** | **Surface sediment**<br>**for microcosm**<br>**establishment** | **Subsurface sediment**<br>**for microcosm**<br>**establishment** | **Hydrocarbon**<br>**assessment** |
| --- | --- | --- | --- | --- | --- |
| 16-06 | 41°37.4476N<br>64°56.3773W | 3260 | 0-20 cmbsf | Not sampled | No faults<br>No acoustic DHIs |
| 16-13 | 41°90.5008N<br>63°47.7986W | 2601 | 0-20 cmbsf | Not sampled | Faults<br>No acoustic DHIs |
| 16-18 | 42°30.8748N<br>62°83.6243W | 2235 | 0-20 cmbsf | 134-141 cmbsf | Faults<br>Acoustic DHIs |
| 16-21 | 42°30.8531N<br>62°83.8306W | 2930 | Not sampled | 142-148 cmbsf | Faults<br>Acoustic DHIs |

<sup>1</sup> All sampling was conducted during the summer in 2016.

### 3.2. Aerobic respiration coupled to hydrocarbon removal at 4℃ in sediment microcosms

Carbon dioxide production and oxygen depletion in the headspace of microcosms were monitored periodically as a proxy for naphtha biodegradation. Figure 3 shows that during the surface sediment incubations, O_2_ depletion and CO_2_ production did not differ significantly between naphtha-amended and unamended control microcosms until after 100 days for 16-13 and 16-18 sediments. On the other hand, sediment 16-06 showed a more rapid response to the naptha amendment (see ANOVA significance values in Table S1). In agreement with O_2_ and CO_2_ observations, GC-MS analysis revealed decreasing concentrations of certain volatile hydrocarbon compounds in the headspace of naphtha-amended microcosms compared to no-sediment controls with identical naphtha amendment (Fig 3G, H & I). However, naphtha-amended microcosms were not substantially different in headspace hydrocarbon profiles compared to heat killed controls (ANOVA values in Table S2).

**Figure 3:**
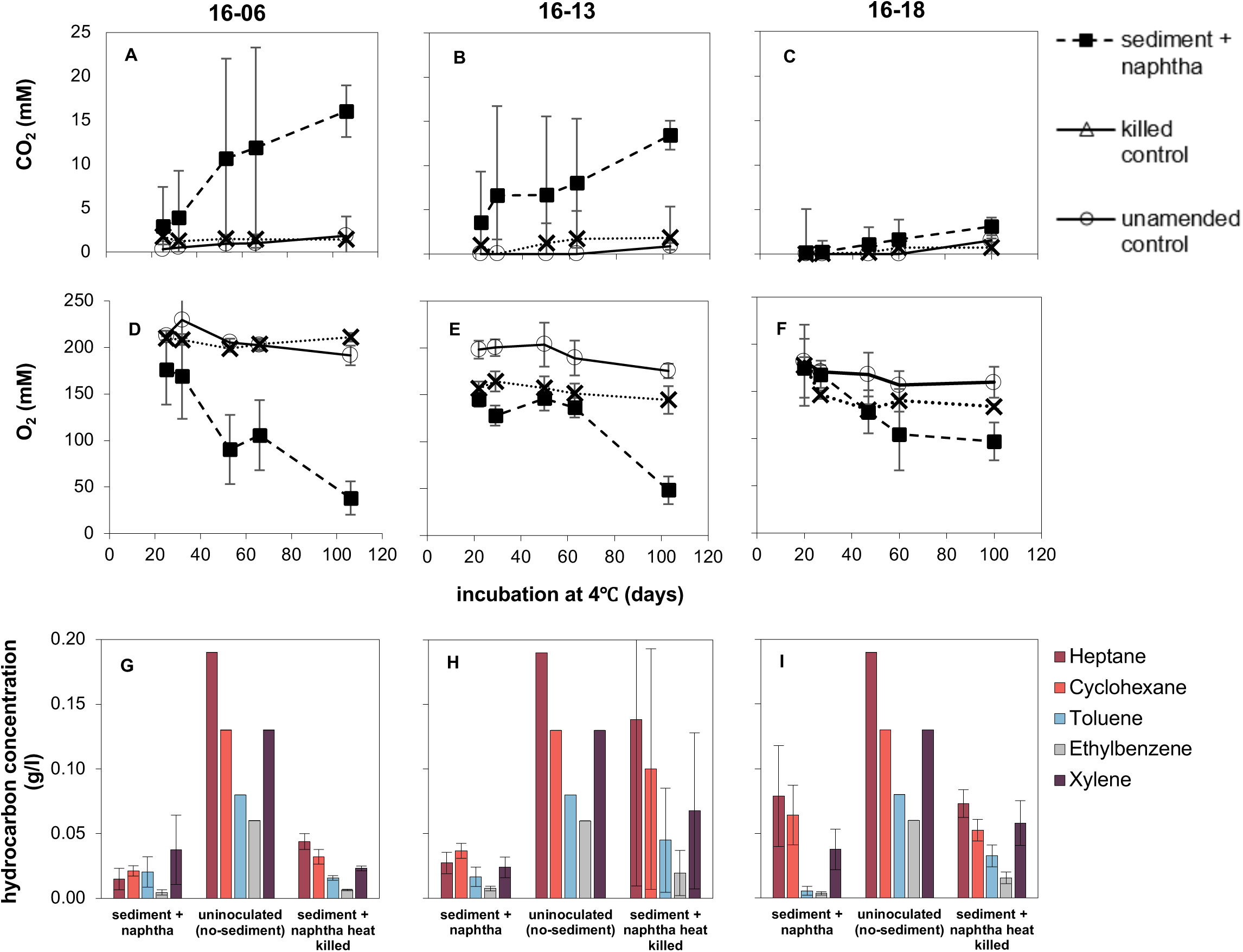
Analysis of carbon dioxide (A-C), oxygen (D-F) and volatile hydrocarbons (G-I) in the headspace of surface sediment (0-20 cmbsf) microcosms during incubation at 4°C. Carbon dioxide production (A-C) and oxygen consumption (D-F) over time reveal most activity during the first 100 days of incubation. Analysis of headspace hydrocarbons after 100 days (G-I) show much lower hydrocarbon concentrations in microcosms that combined sediment and naphtha, compared to uninoculated sediment-free controls (the same data for these controls are plotted beside each of the three sediment-inoculate microcosms in G-I to enable easier comparison). Hydrocarbon concentrations were calculated relative to composition of specific compound in naphtha added.

Deeper anoxic sediment layers exhibited similar potential for aerobic hydrocarbon biodegradation. Naphtha-amended microcosms established with the sediment from the sulfate reduction zone (130 -150 cmbsf) of cores 16-18 and 16-21 gave rise to CO_2_ production that was more extensive than 16-18 surface sediments (0–20 cmbsf), but still not as extensive as the surface sediments from cores 16-06 and 16-13 where hydrocarbons were not detected in situ. Deeper sediment microcosms did not show substantial differences in the depletion of headspace oxygen concentration when compared to controls (Fig. S1).

Hydrocarbon analysis of naphtha-amended microcosms revealed different degrees of hydrocarbon compound depletion relative to heat killed controls between 50 and 100 days of incubation at 4℃. In order to inoculate the microcosms with fresh sediment, incubations were established within hours of sampling on board the ship where headspace measurements of initial volatile hydrocarbon concentrations were not possible; hence the percentage of removal between the period of 50 and 100 days is used for comparison. In general, a greater variety of hydrocarbon compounds were depleted in surface sediment incubations compared to incubation with sediments from deeper anoxic layers (Fig 4; Fig. S2). Toluene, ethylbenzene and xylene were primarily removed from microcosms established with sediment from cores 16-18 and 16-21 where hydrocarbons were detected in situ, whereas microcosms with sediment from core 16-06 showed a depletion of the larger aromatic compounds cyclohexane, ethyl cyclohexane and methyl cyclopentane.

**Figure 4:**
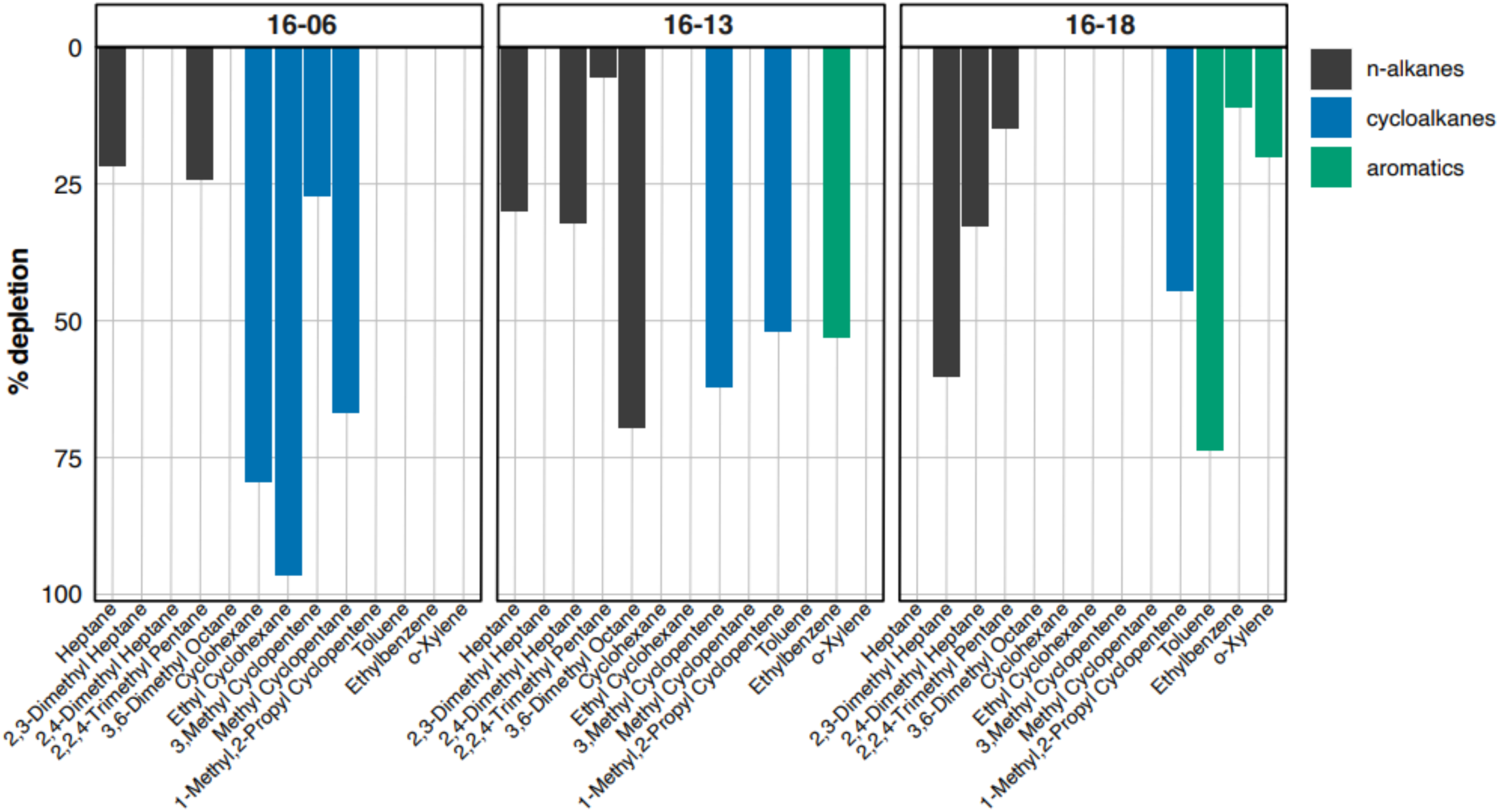
Percentage headspace hydrocarbon depleted in naphtha-amended sediment microcosms relative to heat killed controls for the same sediments between 50 and 100 days of incubation at 4°C.

### 3.3. DNA sequence analysis

Amplicon sequencing resulted in 98 paired end libraries from different microcosms and incubation time points for the various treatments and controls. This resulted in 4,985,354 raw reads (2 × 300 bp), with individual library sizes varying from 1,104 to 542,925 reads (median 63,330 reads). After quality processing and merging of read pairs, the total dataset comprised 2,022,825 reads, with libraries ranging from 3,415 to 245,687 reads (median 24,682 reads) (Table S3). Chimera-free amplicon sequence variants (ASVs) detected by DADA2 differed in length. To avoid spurious ASVs while retaining diversity, all the ASVs shorter than 380 bp or longer than 435 bp were removed. The remaining 14,754 ASVs received taxonomic assignments resulting in 1,195 archaeal, 13,364 bacterial and 91 eukaryotic ASVs as well as 104 ASVs that were unclassified. Corncob identified 29 non-redundant statistically significant ASVs with differential abundance patterns across all experimental parameters that are elaborated on in the next section.

### 3.4. Shifts in microbial community composition during sediment incubations

Comparing microcosms established with sediment from core 16-18 (*in situ* hydrocarbons detected) with the 16-06 and 16-13 microcosms (no hydrocarbons detected in the cores), revealed very few differences (Fig 5). Increases in the relative abundance of *Oleispira* ASV_9, *Pseudomonas* ASV_1 and *Alteromonas* ASV_3 was evident in all surface sediment incubations, compared to being detectable but at low levels (<10%) in parallel incubations without naphtha (Fig. S3). A notable difference was *Colwellia* ASV_6 being significantly elevated in relative abundance only microcosms from cores 16-06 and 16-13, whereas ASV_51 *Moritella* was only detected in microcosms established with 16-06 and 16-18 sediment, albeit at lower relative abundance. Interestingly, the relative abundance of *Oleispira* ASV_9 increased over time in naphtha-amended microcosm from cores 16-06, 16-13, 16-18 but not in parallel unamended microcosms, whereas *Oleispira* ASV_7 exhibited a divergent pattern by increasing in unamended microcosms from core 16-13 sediment (Figure S3) but not in corresponding naphtha-amended microcosms (Fig 5).

**Figure 5:**
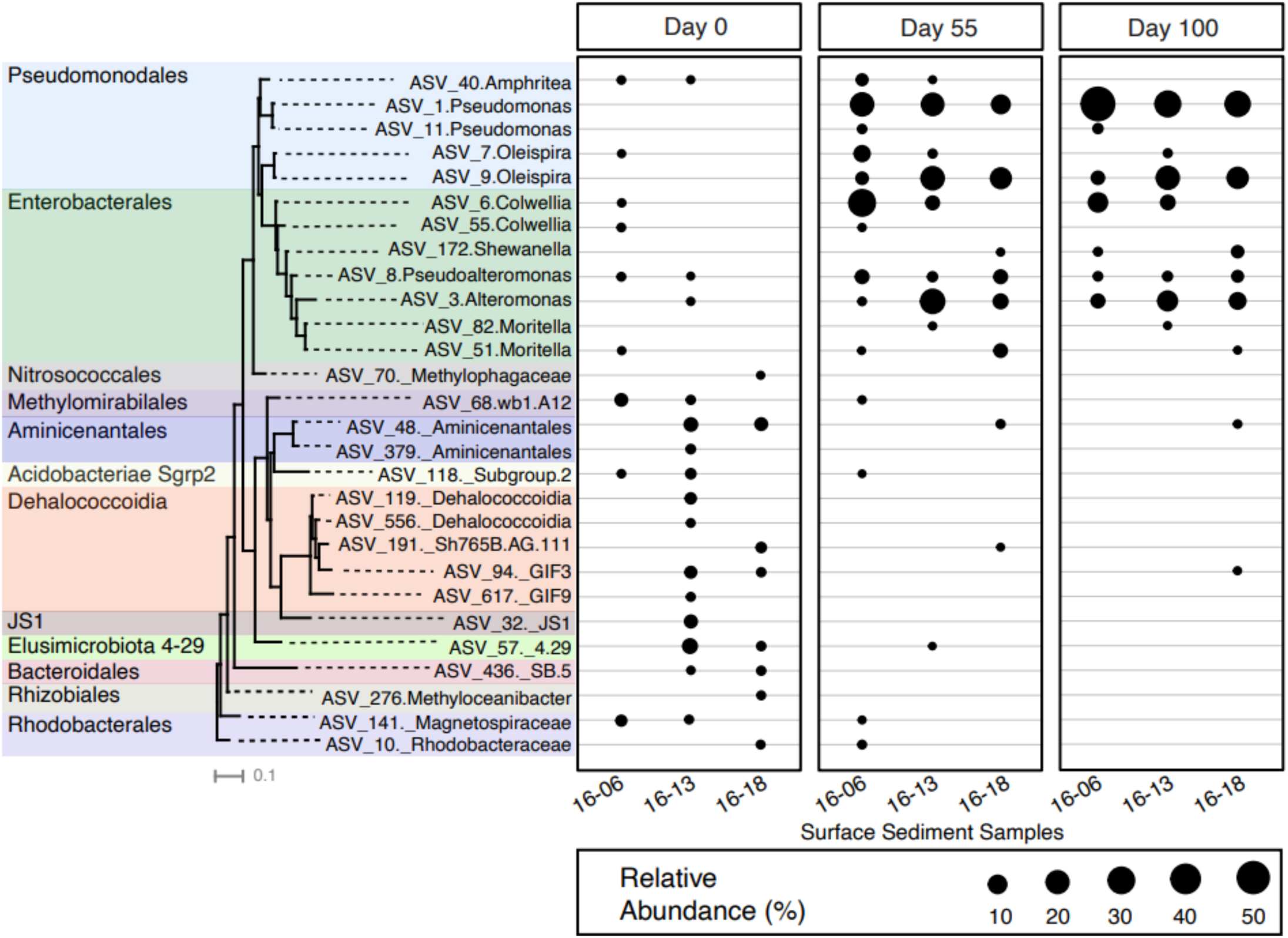
Relative sequence abundance of significantly enriched ASVs in naphtha-amended microcosms over 100 days at 4°C revealed by differential abundance analysis of 16S rRNA gene amplicon libraries. The extra underscore indicates a taxonomy classification at a level higher than genus.

Comparison of microbial communities in oxic naphtha-amended microcosms inoculated with sediment from deeper anoxic layers of cores 16-18 and 16-21 (Fig. S4) revealed general differences relative to the surface sediment incubations. Increased relative abundance of *Pseudomonas* ASV_11 was pronounced in the deeper sediment microcosms (both 16-18 and 16-21). This effect was most dramatic in 16-18 sediment (the only site with parallel incubations of surface and deeper sediment from the same core) where ASV_11 was not enriched in surface sediments but reached >30% in the deeper sediment incubation. In contrast the increase in *Olesipira* ASV_9 in 16-18 surface sediment microcosms was not observed in deeper sediment from the same core (ASV 9 not detected), however this organism did get enriched in the incubation of deeper sediment from core 16-21. The other two populations enriched in all surface sediments, *Pseudomonas* ASV_1 and *Alteromonas* ASV_3, both increased over time in the deeper sediment incubations, where they were similarly among the most prevalent ASVs (Fig. S4).

## 4. Discussion

Oxygen availability is key amidst abiotic and biotic factors that influence hydrocarbon degradation in marine sediments. Microbes preferentially deplete available oxygen during respiration (Breitburg et al., 2010) and as oxygen availability is diminished, alternative electron acceptors, such as sulfate, are used (Lam and Kuypers, 2011). Seabed study sites used here presented not only habitats differing in sulfate concentrations but also proximity to potential hydrocarbon seeps. Additional input of carbon in the form of the hydrocarbon fluids flowing up from below facilitates a more rapid depletion of oxygen leading to the use of sulfate as an alternate electron acceptor at shallower intervals given its high concentration in seawater and marine sediment porewater. The presence of hydrocarbon liquids and gases in cores 16-18 and 16-21 are consistent with the steeper sulfate profiles in these locations. Accordingly, these sites enabled a comparison of biodegradation in deep sea sediments with and without exposure to natural sources of hydrocarbons.

Despite levels of in situ hydrocarbons in cores 16-18 and 16-21 being higher than in cores 16-06 and 16-13 (Figure 2), these hydrocarbon seep sediments did not exhibit a more rapid biodegradation response in 4°C microcosm experiments. Naphtha amendment in surface sediments resulted in enhanced respiration in samples from the top of cores 16-06 and 16-13 relative to surface sediment from core 16-18 where hydrocarbons were detected (Figure 3). Compared to controls incubated in parallel without hydrocarbon amendment, significantly higher carbon dioxide production (P = 0.03) and oxygen depletion (P = 0.001) were recorded in 0006 sediment amended with naphtha after 60 days. This rapid <60-day response was not observed in naphtha-amended surface sediments from site 16-13, but by 100 days activity in the 16-13 microcosms was comparable to the 0006 microcosms, i.e., >10 mM CO_2_ production, whereas only 3 mM CO_2_ production was measured in the naphtha-amended incubation of surface sediment from core 0018 (Fig 3; Fig S1; Table S1). These experiments do not provide evidence to suggest that prior exposure to hydrocarbons due to seepage leads to an enhanced biodegradation response.

Enrichment of Gammaproteobacterial groups in all microcosms reflects observations in other marine systems in response to oil spills. Gammaproteobacteria were not among the taxa deemed to be diagnostic for thermogenic hydrocarbon seepage in the deep-sea sediments at this study site (Li et al. 2023) or at Gulf of Mexico seabed hydrocarbon seeps (Chakraborty et al. 2020) where lineages including *Caldatribacteria*, *Aminicenantes* and *Campilobacterota* are common indicators. Instead, naphtha resulted in the enrichment of *Oleispira*, *Alteromonas* and *Pseudomonas* as being among the most important populations in the biodegradation response. Some of these gammaproteobacterial groups, such as *Oleispira*, are considered obligate hydrocarbonoclastic taxa (Yakimov et al., 2007) that are otherwise minor bacterial constituents of the pristine (oil-free) marine systems (Radwan et al., 2019). Different *Oleispira* strains were enriched in both naphtha amended (ASV_9) and unamended (ASV_7) 4°C incubations over the course of 100 days (Fig. 5 and S3), calling into question the designation of this genus as being ‘obligately’ hydrocarbonoclastic. Non-detection of *Oleispira* ASV_9 at the onset of the incubations (week 0) in all but one of the sediments (i.e., the deeper anoxic layer of core 16-21; Fig. S3) is consistent with hydrocarbon-degrading taxa being minor constituents with low in situ abundance. On the other hand, *Alteromonas* ASV_3, which was enriched in the presence of naphtha in all five microcosms spanning surface and deeper sediments, could be detected in three out of the five unincubated in situ sediment samples (i.e., week 0) at 0.04 to 0.17% relative sequence abundance. This is consistent with its 16S rRNA gene phylogeny indicating a close relationship with the known hydrocarbon degrader *Alteromonas naphthalenivorans* (Fig. S5).

Instances of *Alteromonas* detection in situ include both 16-18 and 16-21 sediments where hydrocarbons were detected in the sediment cores (Fig. 1). Detection and enrichment of *Alteromonas*, *Pseudomonas* and other aerobic hydrocarbon degrading Gammaproteobacteria both in surface sediment and down to 148 cmbsf in these two locations (Figure 1A, B) suggests that these aerobic hydrocarbon degraders persist in a viable state during sediment burial to this depth, which lasted thousands of years in this area given the low sedimentation rate of 0.4 mm y^-1^ (Normandeau & Campbell, 2020).

Metagenomic and metatranscriptomic analyses conducted during the unmitigated release of oil for 84 days during DWH blowout revealed alkane degradation was a dominant hydrocarbon-degrading pathway coinciding with Gammaproteobacteria including *Pseudomonas* being dominant members of the community (Mason et al., 2012). In our study, an increased relative abundance of different *Pseudomonas* ASVs was observed in oxic naphtha-amended microcosms, corresponding to headspace depletion of alkanes and aromatic compounds, as has been reported for other members of this genus (Whyte et al., 1997). Toluene removal was especially prevalent in the 16-18 and 16-21 microcosms during 50-100 days, corresponding with increased abundance of *Pseudomonas* ASVs (Fig.4; Fig. S2). More rapid hydrocarbon metabolism may be explained by the observed natural hydrocarbon seepage at this location (Figure 1B). This could be relevant in the context of deep marine ecosystems like this study site and the DWH location in the Gulf of Mexico where there will be reduced weathering of light hydrocarbons in released crude oil due to the cold seawater with resultant higher concentrations of volatile toxic compounds such as BTEX (Brakstad et al., 2017). Widespread capacity for alkane biodegradation in the marine microbiome may not depend solely on the presence of natural hydrocarbon seeps. Alkane production by marine phototrophs exerts more widespread selective pressure for alkane biodegradation (Lea-Smith et al. 2015; Love et al. 2021). For aromatic hydrocarbons, seabed seeps may be a more likely point of introduction into the marine environment, resulting in ‘priming’ the marine microbiome for aromatic metabolism. Despite the overall slower response to naphtha amendment in the seep sediments, as described above, aromatic hydrocarbons were more readily consumed by the microbial communities enriched from the 16-18 and 16-21 sediments (Fig. 4; Fig. S3) where there is prior exposure to thermogenic hydrocarbons (Figs. 1, 2). *Pseudomonas* spp. are well documented for their ability to degrade BTEX (Chicca et al., 2020) with the *Pseudomonas* ASVs in this study being closely related to those in other hydrocarbon-degrading systems (Fig. S5). For example, enrichment of *Alteromonas* in similar cold deep sea enrichment cultures amended with crude oil has also been linked to the degradation of aromatic compounds (Cui et al. 2008).

Permanently cold deep-sea sediments and their incubation with hydrocarbons at 4°C also expand knowledge of marine biodegradation processes and populations that operate at low temperatures. While *Oleispira*, *Alteromonas*, *Pseudomonas* and other ‘usual suspects’ within the Gammaproteobacteria are typically considered mesophiles, it is inferred from their enrichment here and their close phylogenetic relationships to bacteria from cold environments (Fig. S5) that these strains are psychrophilic inhabitants of the deep sea. Many psychrotolerant and some psychrophilic strains of *Pseudomonas* have been isolated (Canion et al., 2013; Kim et al., 2013; Kosina et al., 2013). Other studies have similarly shown that *Pseudomonas* spp. are found in low abundance across a range of cold environments and become dominant under stress, such as acute hydrocarbon exposure (Farrell et al., 2003; Aislabe et al., 2006; Yergeau et al., 2012). Enrichment of *Oleispira* ASVs in naphtha-amended microcosms is similarly unsurprising. Gregson et al. (2020) highlight that *Oleispira* shares many traits with other described genera of well-known marine obligate hydrocarbon degraders like *Alcanivorax* (Yakimov *et al*., 1998) and *Thalassolituus* (Yakimov *et al*., 2004) including marine origin, respiratory metabolism and ability to metabolise aliphatic alkanes and their derivatives. However, in contrast to other so-called obligate genera, *Oleispira antarctica* was shown to exhibit a broad growth temperature optimum between 1°C and 15°C (Yakimov *et al*., 2003) suggesting a potential for *Oleispira* spp. to dominate microbial communities due to their ecological competitiveness in cold environments (Hazen *et al*., 2010; Mason *et al*., 2012; Kube et al., 2013).

## 5. Conclusions

Microcosm experiments with deep sea sediments sampled from sites with and without background thermogenic hydrocarbon seepage did not support the hypothesis that prior exposure to hydrocarbons would lead to enhanced biodegradation. The capacity for the biodegradation of spilled petroleum compounds by marine microbial communities is often explained by the presence of natural hydrocarbon seeps in the seabed. Chemosynthetic ecosystems fuelled by thermogenic hydrocarbons highlight an important role for microbial populations capable of oxidizing these compounds as primary producers in the seabed (Dong et al 2019; 2020). Despite this premise, bacterial groups like *Oleispira*, *Alteromonas*, *Pseudomonas* and other members of the Gammaproteobacteria that typically respond to oil spills or oil-amendment enrichment experiments, like the ones performed here, differ from the signature microbial groups that define cold seep sediments such as *Caldatribacteria*, *Aminicenantes* and *Campilobacterota*. In the present study, none of the latter groups became enriched when sediments were exposed to low molecular weight hydrocarbons (naphtha) over a 100-day period. Instead, relatively rare psychrophilic Gammaproteobacteria responded to acute hydrocarbon exposures, supporting and expanding the widespread importance of these aerobic hydrocarbon-degrading bacteria across a range of marine oil pollution scenarios.

## Credit author statement

Oyeboade Adebayo – Conceptualization; Formal analysis; Investigation; Methodology; Writing-Original Draft; Writing-Review & Editing; Visualization

Srijak Bhatnagar – Data curation; Formal analysis, Writing-Review & Editing, Visualization

Jamie Webb – Formal analysis

Calvin Campbell – Project administration; Resources; Supervision

Martin Fowler – Formal analysis; Writing-Review & Editing

Natasha M. MacAdam – Formal analysis

Adam MacDonald – Project administration; Resources; Supervision

Carmen Li – Investigation

Casey R.J. Hubert – Conceptualization; Funding acquisition; Supervision; Writing-Review & Editing

## Acknowledgements

This research was supported by funds from the Campus Alberta Innovates Program (CAIP) chair program, Genome Canada, and the Marine Environment Observation Prediction and Response (MEOPAR) network. Partnerships with Genome Atlantic, Genome Alberta, the Government of Nova Scotia and Natural Resources Canada are gratefully acknowledged. Sampling aboard CCGS *Hudson* was made possible by its captain and crew, and collaborative support of Natural Resources Canada. Project management and support were provided by Dr Rhonda Clark and Carey Ryan.

## Declaration of competing interest

Authors declare that they have no known competing financial interests or personal relationships that could influence the work reported in this paper.

**Appendix: Supplementary data**

---

**Figure S1:**
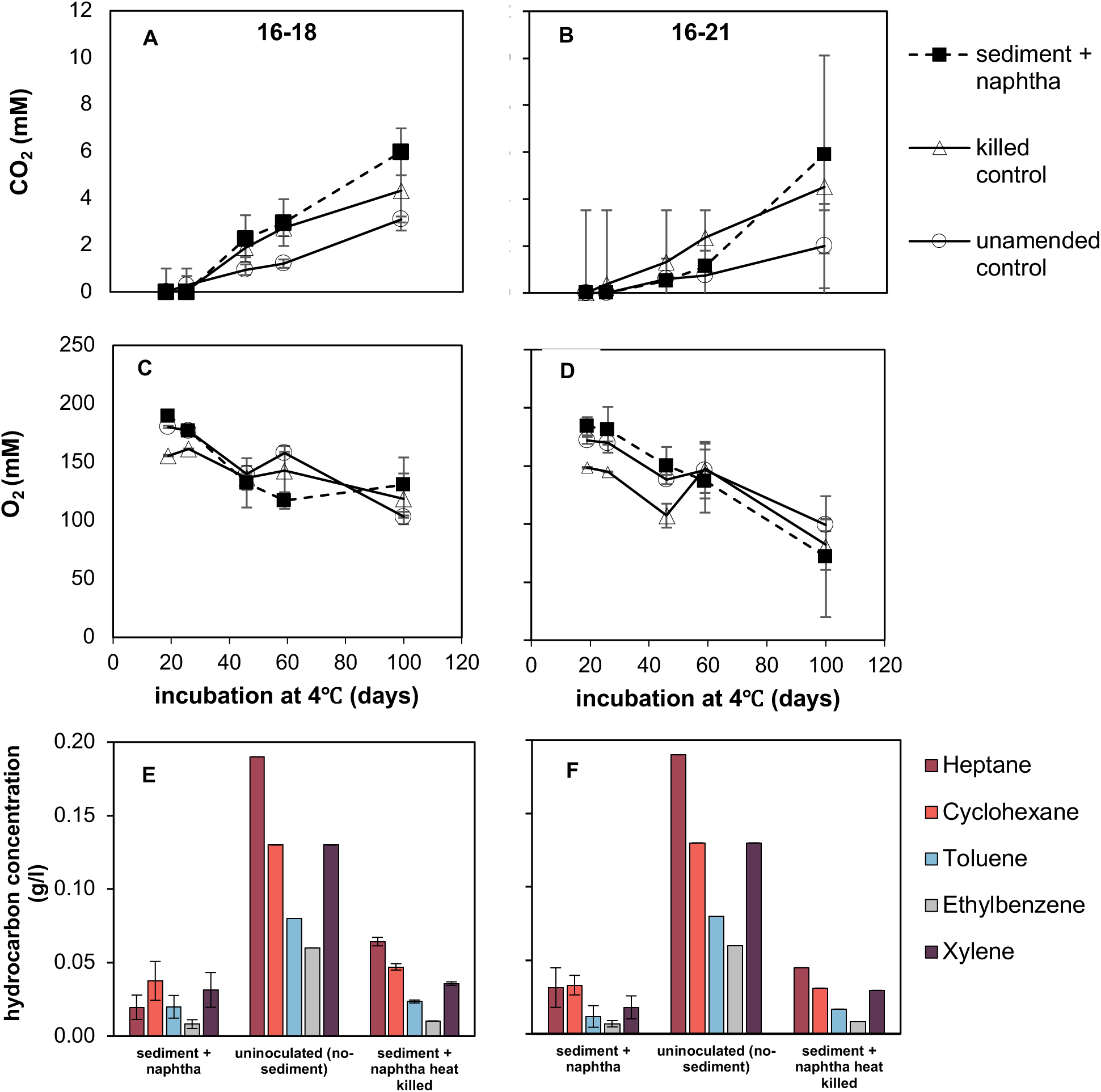
Analysis of carbon dioxide (A, B), oxygen (C, D) and volatile hydrocarbons (E, F) in the headspace of deeper sediment 16-18 (134-141 cmbsf), 16-21 (142-148 cmbsf) microcosms during incubation at 4°C. Carbon dioxide production (A, B) over time reveal most activity during the first 100 days of ncubation. Analysis of headspace hydrocarbons after 100 days (E, F) show much ower hydrocarbon concentrations in microcosms that combined sediment and naphtha, compared to uninoculated sediment-free controls (the same data for hese controls are plotted beside each of the two sediment-inoculate microcosms in E & F to enable easier comparison). Hydrocarbon concentrations were calculated elative to composition of specific compound in naphtha added.

**Figure S2:**
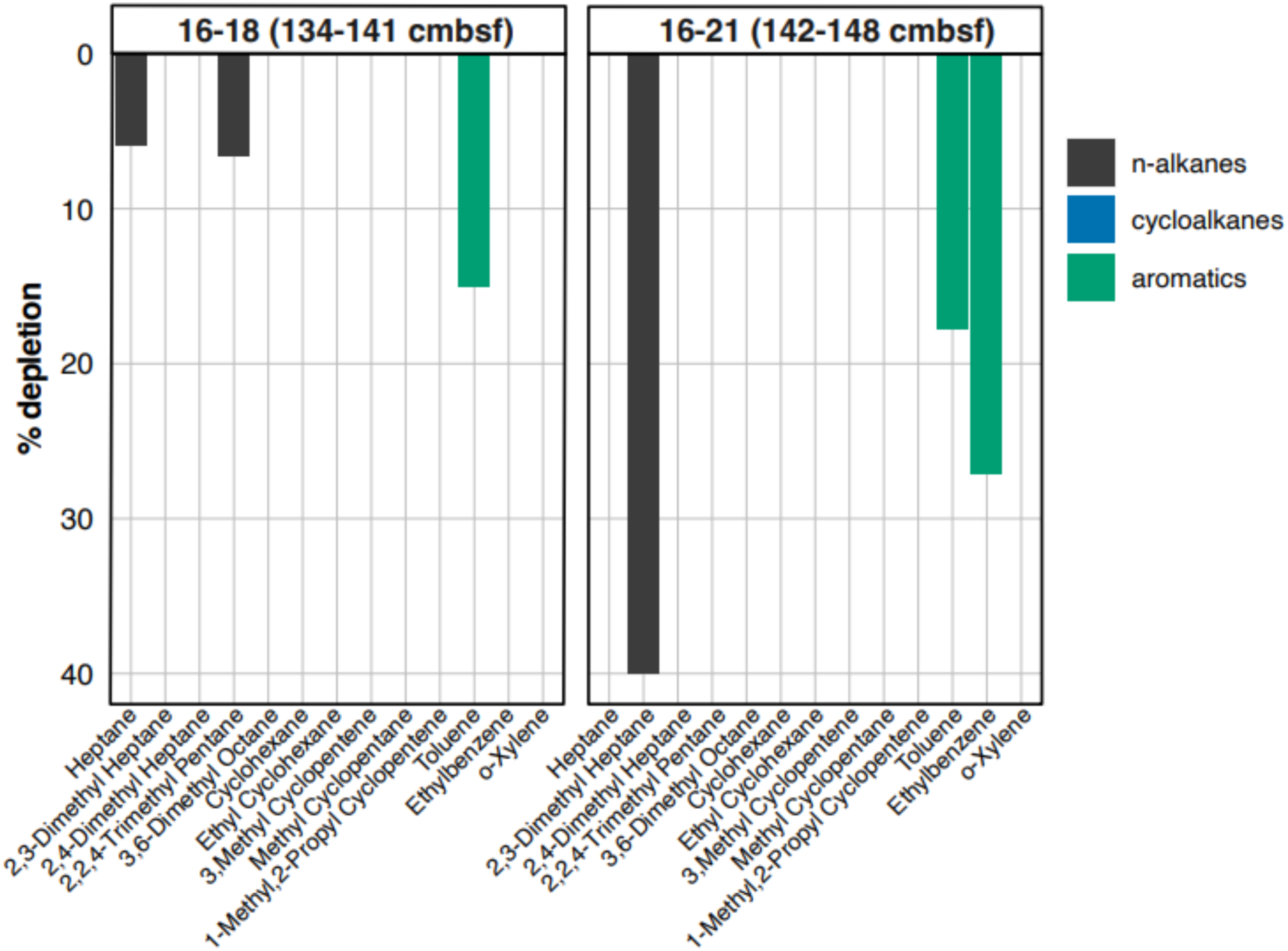
Percentage headspace hydrocarbon depleted in naphtha amended subsurface sediment microcosms relative to heat killed controls for the same sediments between 50 and 100 days of incubation at 4°C.

**Figure S3:**
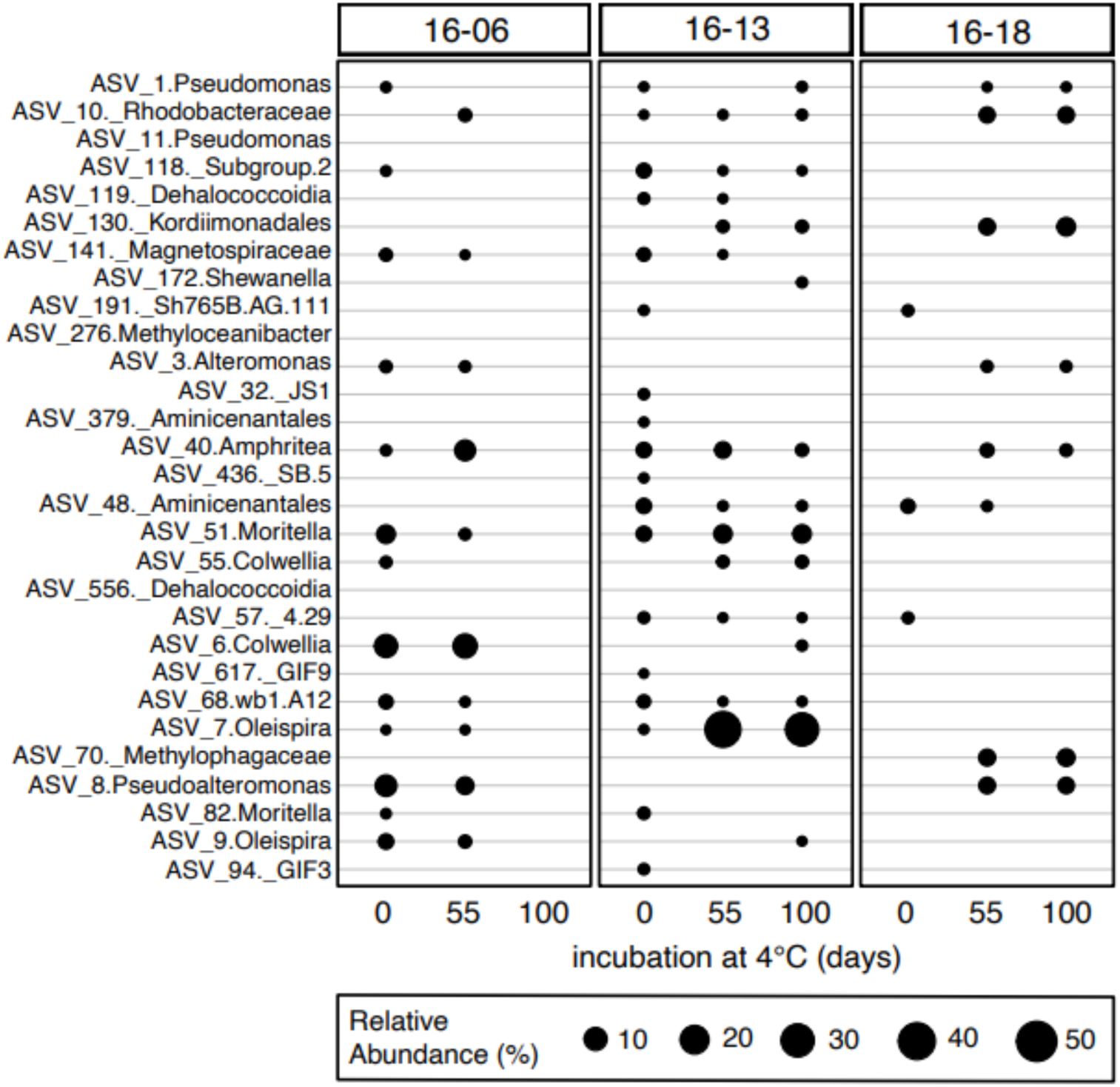
Relative sequence abundance of statistically significant ASVs revealed by differential analysis based on 16S rRNA gene amplicons showing community change in unamended microcosms inoculated with surface sediment during a period of 100 days incubation at 4°C. The extra underscore indicates a taxonomy classification at a level higher than genus.

**Figure S4:**
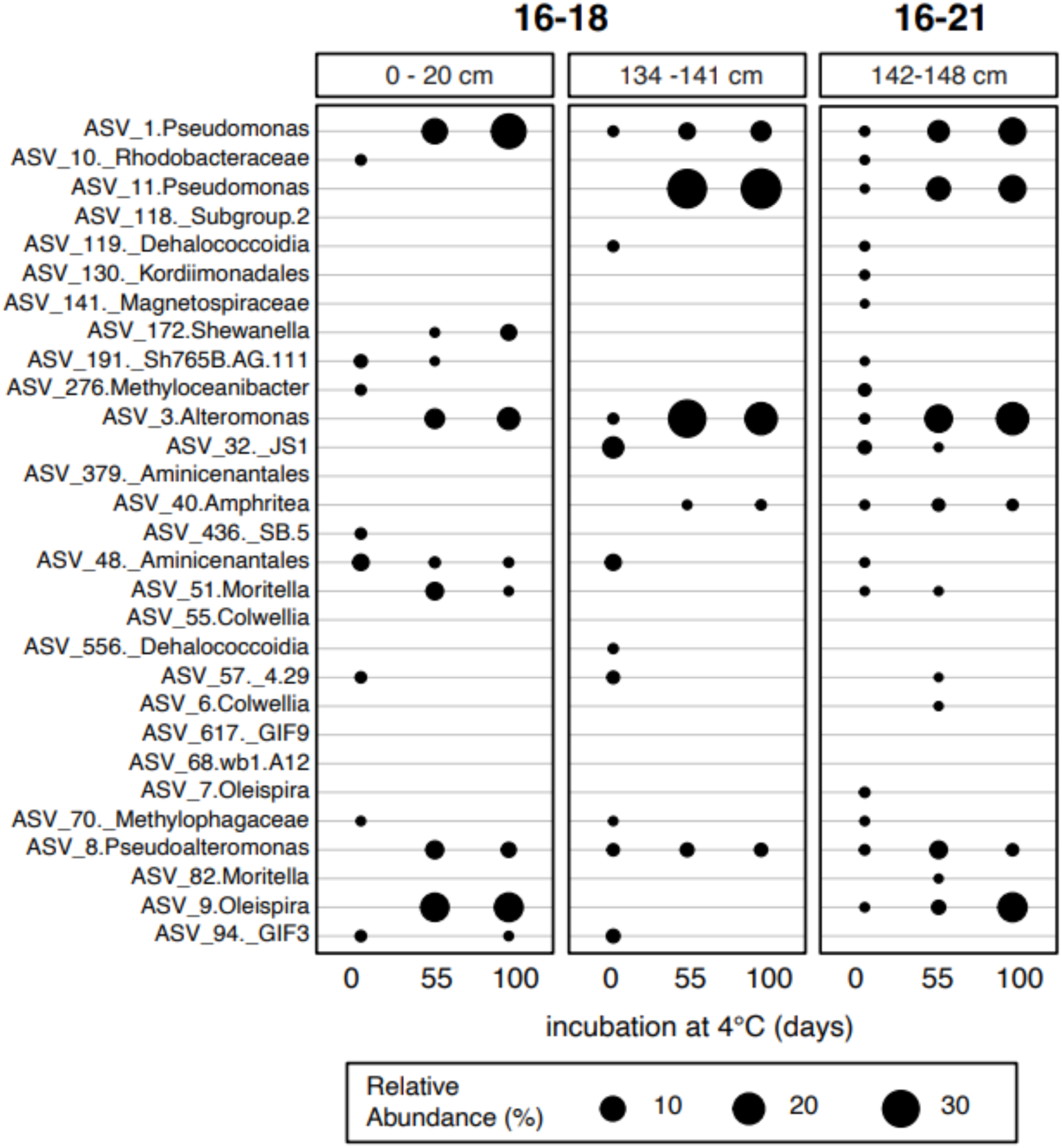
Relative sequence abundance of statistically significant ASVs revealed by differential analysis based on 16S rRNA gene amplicons showing community change in naphtha-amended microcosms inoculated with surface and deeper sediment (16-18) and deeper sediment (16-21) during a period of 100 days incubation at 4°C. The extra underscore indicates a taxonomy classification at a level higher than genus.

**Figure S5:**
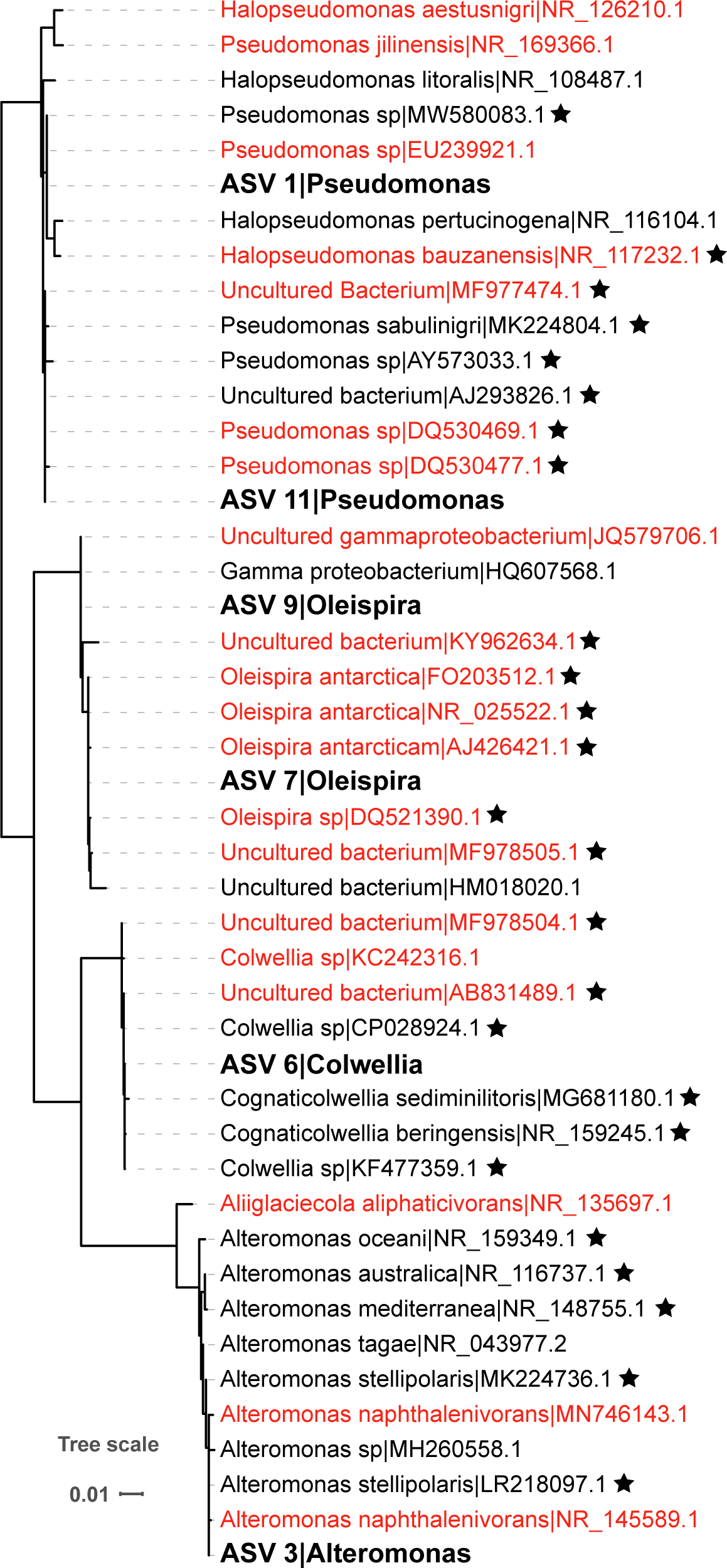
Phylogeny of ASVs from sediment incubations (bold) affiliated with known hydrocarbon-degrading genera and their close sequence matches (≥99% sequence identity) in the NCBI nr database. NCBI hits annotated as being from hydrocarbon-associated environments are highlighted in red, and hits with cold tolerance as demonstrated through their cultivation or inferred from the environment are marked by a star.

**Table S1.** Analysis of variance (ANOVA) comparing respiration in naphtha amendment, heat killed and unamended microcosms of each tested core.

| Core | Day |  | Sum Sq | Mean Sq | F Val | Probability |
| --- | --- | --- | --- | --- | --- | --- |
| 16 - 06<br>(0-20cm) | 20 | O<sub>2</sub> | 2531 | 1265.5 | 26.67 | 0.15 |
|  | 60 |  | 25039 | 12520 | 26.63 | 1.04 e-03** |
|  | 100 |  | 53808 | 26904 | 177 | 6e-06*** |
|  | 20 | CO<sub>2</sub> | 31.6 | 15.80 | 2.32 | 0.18 |
|  | 60 |  | 530.0 | 265.02 | 6.16 | 0.0351* |
|  | 100 |  | 1080.3 | 540.2 | 186.7 | 3.96e-06*** |
| 16 - 13<br>(0-20cm) | 20 | O<sub>2</sub> | 5483 | 2742 | 1.933 | 0.225 |
|  | 60 |  | 5182 | 2591 | 1.786 | 0.246 |
|  | 100 |  | 30758 | 15379 | 35.29 | 4.81 e-04*** |
|  | 20 | CO<sub>2</sub> | 23.86 | 11.93 | 0.933 | 0.444 |
|  | 60 |  | 124.0 | 61.98 | 2.996 | 0.125 |
|  | 100 |  | 370.6 | 185.31 | 178.6 | 4.51e-06*** |
| 16 - 18<br>(0-20cm) | 20 | O<sub>2</sub> | 79 | 39.6 | 0.042 | 0.959 |
|  | 60 |  | 4320 | 2159.8 | 3.58 | 0.0948 |
|  | 100 |  | 5971 | 2985.6 | 13.18 | 6.38 e-03** |
|  | 20 | CO<sub>2</sub> | 0.0857 | 0.0428 | 0.99 | 0.422 |
|  | 60 |  | 4.141 | 2.071 | 1.276 | 0.345 |
|  | 100 |  | 8.749 | 4.375 | 12.5 | 7.25 e-03** |
| 16 - 18<br>(134-141cm) | 20 | O<sub>2</sub> | 1231.2 | 615.6 | 84.63 | 0.0023** |
|  | 60 |  | 1692 | 845.8 | 2.323 | 0.246 |
|  | 100 |  | 750.5 | 375.5 | 1.085 | 0.442 |
|  | 20 | CO<sub>2</sub> | NA | NA | NA | NA |
|  | 60 |  | 3.622 | 1.8112 | 4.113 | 0.138 |
|  | 100 |  | 8.401 | 4.201 | 2.612 | 0.22 |
| 16 - 21<br>(142-148 cm) | 20 | O<sub>2</sub> | 1532 | 766 | 16.93 | 0.0112* |
|  | 60 |  | 179.9 | 90 | 0.243 | 0.795 |
|  | 100 |  | 892 | 445.9 | 0.322 | 0.742 |
|  | 20 | CO<sub>2</sub> | NA | NA | NA | NA |
|  | 60 |  | 2.858 | 1.429 | 5.067 | 0.080 |
|  | 100 |  | 18.34 | 9.170 | 1.716 | 0.29 |

Sum Sq: Sum of squares; Mean Sq: Mean sum of squares.
F val: F value; Signif. codes: \*\*\* P 0.001 \*\* P 0.01 \* P 0.05
NA: Not applicable as all values were 0.

**Table S2.** Analysis of variance (ANOVA) comparing hydrocarbon concentration in naphtha amendment, heat killed and unamended microcosms of each tested core after 100 days of incubation at 4℃.

| Core | Hydrocarbon | Sum Sq | Mean Sq | F Val | Probability |
| --- | --- | --- | --- | --- | --- |
| 16 - 06<br>(0-20cm) | Heptane | 0.02 | 0.01 | 221.19 | 8.03 e -05*** |
|  | Cyclohexane | 0.001 | 0.005 | 195.59 | 1.02 e-04*** |
|  | Toluene | 0.003 | 0.001 | 23.61 | 6.09 e-03** |
|  | Ethylbenzene | 0.003 | 0.001 | 570.67 | 1.22 e-05*** |
|  | Xylene | 0.02 | 0.01 | 41.99 | 0.02* |
| 16 - 13<br>(0-20cm) | Heptane | 0.03 | 0.01 | 1.70 | 0.29 |
|  | Cyclohexane | 0.01 | 0.004 | 1.07 | 0.42 |
|  | Toluene | 0.003 | 0.002 | 1.95 | 0.26 |
|  | Ethylbenzene | 0.002 | 0.001 | 6.69 | 0.05* |
|  | Xylene | 0.009 | 0.004 | 2.41 | 0.2 |
| 16 - 18<br>(0-20cm) | Heptane | 0.01 | 0.006 | 0.79 | 0.51 |
|  | Cyclohexane | 0.005 | 0.002 | 0.89 | 0.47 |
|  | Toluene | 0.004 | 0.002 | 6.51 | 0.06 |
|  | Ethylbenzene | 0.002 | 0.001 | 12.87 | 0.02* |
|  | Xylene | 0.006 | 0.003 | 1.41 | 0.34 |
| 16 - 18<br>(134-141cm) | Heptane | 0.019 | 0.009 | 252.93 | 3.93 e-03** |
|  | Cyclohexane | 0.006 | 0.003 | 34.79 | 0.03* |
|  | Toluene | 0.003 | 0.001 | 45.12 | 0.02* |
|  | Ethylbenzene | 0.002 | 0.001 | 231.05 | 4.30 e-03** |
|  | Xylene | 0.007 | 0.004 | 52.68 | 0.02* |
| 16 - 21<br>(142-148 cm) | Heptane | 0.019 | 0.009 | 53.53 | 0.02* |
|  | Cyclohexane | 0.008 | 0.004 | 85.61 | 0.01* |
|  | Toluene | 0.004 | 0.002 | 33.37 | 0.03* |
|  | Ethylbenzene | 0.002 | 0.001 | 230.98 | 4.31 e-03** |
|  | Xylene | 0.009 | 0.005 | 79.09 | 0.01* |

Sum Sq: Sum of squares; Mean Sq: Mean sum of squares.
F val: F value; Signif. codes: \*\*\* P 0.001 \*\* P 0.01 \* P 0.05

**Table S3.** Description and read count for 16S rRNA gene amplicon libraries.

| **Library** | **Core** | **Treatment** | **Core Section** | **Time** | **Rep** | **Reads-**<br>**Raw** | **Reads-**<br>**Filtered** |
| --- | --- | --- | --- | --- | --- | --- | --- |
| 13_PC_BS_0cm_TBS_1 | 16-13 | in situ | 0cm | TBS | 1 | 68913 | 28554 |
| 13_PC_BS_0cm_TBS_2 | 16-13 | in situ | 0cm | TBS | 2 | 28013 | 13378 |
| 13_PC_BS_0cm_TBS_3 | 16-13 | in situ | 0cm | TBS | 3 | 16197 | 6778 |
| 13_PC_BS_20cm_TBS_1 | 16-13 | in situ | 20cm | TBS | 1 | 37582 | 14580 |
| 13_PC_BS_20cm_TBS_2 | 16-13 | in situ | 20cm | TBS | 2 | 25532 | 12436 |
| 13_PC_BS_20cm_TBS_3 | 16-13 | in situ | 20cm | TBS | 3 | 8901 | 3415 |
| 13_PC_NA_OX_T0_1 | 16-13 | Naphtha | 0-20 | Day 0 | 1 | 63427 | 31508 |
| 13_PC_NA_OX_T0_2 | 16-13 | Naphtha | 0-20 | Day 0 | 2 | 50127 | 24589 |
| 13_PC_NA_OX_T0_3 | 16-13 | Naphtha | 0-20 | Day 0 | 3 | 120670 | 58376 |
| 13_PC_NA_OX_T1_1 | 16-13 | Naphtha | 0-20 | Day 50 | 1 | 100964 | 53304 |
| 13_PC_NA_OX_T1_2 | 16-13 | Naphtha | 0-20 | Day 50 | 2 | 84322 | 56213 |
| 13_PC_NA_OX_T1_3 | 16-13 | Naphtha | 0-20 | Day 50 | 3 | 28467 | 14538 |
| 13_PC_NA_OX_T2_1 | 16-13 | Naphtha | 0-20 | Day 100 | 1 | 26350 | 13251 |
| 13_PC_NA_OX_T2_2 | 16-13 | Naphtha | 0-20 | Day 100 | 2 | 70306 | 47777 |
| 13_PC_NA_OX_T2_3 | 16-13 | Naphtha | 0-20 | Day 100 | 3 | 67189 | 34131 |
| 13_PC_UN_OX_T0_3 | 16-13 | Naphtha | 0-20 | Day 0 | 3 | 77685 | 35965 |
| 13_PC_UN_OX_T1_3 | 16-13 | Naphtha | 0-20 | Day 50 | 3 | 49951 | 28427 |
| 13_PC_UN_OX_T2_3 | 16-13 | Naphtha | 0-20 | Day 100 | 3 | 90644 | 53020 |
| 18_PC_NA_AN_T0_1 | 16-18 | Naphtha | 134-141 | Day 0 | 1 | 102873 | 21871 |
| 18_PC_NA_AN_T0_2 | 16-18 | Naphtha | 134-141 | Day 0 | 2 | 37588 | 5618 |
| 18_PC_NA_AN_T1_1 | 16-18 | Naphtha | 134-141 | Day 50 | 1 | 52960 | 12134 |
| 18_PC_NA_AN_T1_2 | 16-18 | Naphtha | 134-141 | Day 50 | 2 | 55584 | 11068 |
| 18_PC_NA_AN_T2_1 | 16-18 | Naphtha | 134-141 | Day 100 | 1 | 61778 | 12414 |
| 18_PC_NA_AN_T2_2 | 16-18 | Naphtha | 134-141 | Day 100 | 2 | 48266 | 11159 |
| 18_PC_NA_OX_T0_1 | 16-18 | Naphtha | 0-20 | Day 0 | 1 | 53631 | 9050 |
| 18_PC_NA_OX_T0_2 | 16-18 | Naphtha | 0-20 | Day 0 | 2 | 126587 | 24111 |
| 18_PC_NA_OX_T0_3 | 16-18 | Naphtha | 0-20 | Day 0 | 3 | 45988 | 7220 |
| 18_PC_NA_OX_T1_1 | 16-18 | Naphtha | 0-20 | Day 50 | 1 | 59846 | 12285 |
| 18_PC_NA_OX_T1_2 | 16-18 | Naphtha | 0-20 | Day 50 | 2 | 73274 | 9275 |
| 18_PC_NA_OX_T1_3 | 16-18 | Naphtha | 0-20 | Day 50 | 3 | 80382 | 17898 |
| 18_PC_NA_OX_T2_1 | 16-18 | Naphtha | 0-20 | Day 100 | 1 | 46501 | 10074 |
| 18_PC_NA_OX_T2_2 | 16-18 | Naphtha | 0-20 | Day 100 | 2 | 37888 | 7957 |
| 18_PC_NA_OX_T2_3 | 16-18 | Naphtha | 0-20 | Day 100 | 3 | 48739 | 8717 |
| 18_PC_UN_OX_T0_3 | 16-18 | Unamended | 0-20 | Day 0 | 3 | 42398 | 8059 |
| 18_PC_UN_OX_T1_3 | 16-18 | Unamended | 0-20 | Day 50 | 3 | 63664 | 15785 |
| 18_PC_UN_OX_T2_3 | 16-18 | Unamended | 0-20 | Day 100 | 3 | 84820 | 22177 |
| 21_PC_NA_AN_T0_1 | 16-21 | Naphtha | 142-148 | Day 0 | 1 | 542925 | 245687 |
| 21_PC_NA_AN_T0_2 | 16-21 | Naphtha | 142-148 | Day 0 | 2 | 53654 | 24015 |
| 21_PC_NA_AN_T0_3 | 16-21 | Naphtha | 142-148 | Day 0 | 3 | 74551 | 32453 |
| 21_PC_NA_AN_T1_1 | 16-21 | Naphtha | 142-148 | Day 50 | 1 | 111523 | 52267 |
| 21_PC_NA_AN_T1_2 | 16-21 | Naphtha | 142-148 | Day 50 | 2 | 85445 | 35070 |
| 21_PC_NA_AN_T1_3 | 16-21 | Naphtha | 142-148 | Day 50 | 3 | 99668 | 43013 |
| 21_PC_NA_AN_T2_1 | 16-21 | Naphtha | 142-148 | Day 100 | 1 | 87234 | 38298 |
| 21_PC_NA_AN_T2_2 | 16-21 | Naphtha | 142-148 | Day 100 | 2 | 105263 | 43592 |
| 21_PC_NA_AN_T2_3 | 16-21 | Naphtha | 142-148 | Day 100 | 3 | 63234 | 26824 |
| 21_PC_UN_AN_T0_1 | 16-21 | Unamended | 142-148 | Day 0 | 1 | 1104 | 145 |
| 21_PC_UN_AN_T0_2 | 16-21 | Unamended | 142-148 | Day 0 | 2 | 52026 | 23598 |
| 21_PC_UN_AN_T1_1 | 16-21 | Unamended | 142-148 | Day 50 | 1 | 116479 | 54463 |
| 21_PC_UN_AN_T1_2 | 16-21 | Unamended | 142-148 | Day 50 | 2 | 82790 | 36920 |
| 21_PC_UN_AN_T2_1 | 16-21 | Unamended | 142-148 | Day 100 | 1 | 27585 | 10381 |
| 21_PC_UN_AN_T2_2 | 16-21 | Unamended | 142-148 | Day 100 | 2 | 73325 | 29910 |
| 6_PC_BS_0cm_TBS_1 | 16-06 | in situ | 0cm | TBS | 1 | 107353 | 37578 |
| 6_PC_BS_0cm_TBS_2 | 16-06 | in situ | 0cm | TBS | 2 | 86573 | 31704 |
| 6_PC_BS_0cm_TBS_3 | 16-06 | in situ | 0cm | TBS | 3 | 68553 | 25839 |
| 6_PC_BS_20cm_TBS_1 | 16-06 | in situ | 20cm | TBS | 1 | 61646 | 21459 |
| 6_PC_BS_20cm_TBS_2 | 16-06 | in situ | 20cm | TBS | 2 | 102366 | 38911 |
| 6_PC_BS_20cm_TBS_3 | 16-06 | in situ | 20cm | TBS | 3 | 56663 | 19880 |
| 6_PC_NA_OX_T0_1 | 16-06 | Naphtha | 0-20 | Day 0 | 1 | 56220 | 24682 |
| 6_PC_NA_OX_T0_2 | 16-06 | Naphtha | 0-20 | Day 0 | 2 | 52597 | 22262 |
| 6_PC_NA_OX_T0_3 | 16-06 | Naphtha | 0-20 | Day 0 | 3 | 103443 | 42542 |
| 6_PC_NA_OX_T1_1 | 16-06 | Naphtha | 0-20 | Day 50 | 1 | 99888 | 56890 |
| 6_PC_NA_OX_T1_2 | 16-06 | Naphtha | 0-20 | Day 50 | 2 | 96378 | 55388 |
| 6_PC_NA_OX_T1_3 | 16-06 | Naphtha | 0-20 | Day 50 | 3 | 54027 | 29668 |
| 6_PC_NA_OX_T2_1 | 16-06 | Naphtha | 0-20 | Day 100 | 1 | 44976 | 23396 |
| 6_PC_NA_OX_T2_2 | 16-06 | Naphtha | 0-20 | Day 100 | 2 | 67414 | 33963 |
| 6_PC_NA_OX_T2_3 | 16-06 | Naphtha | 0-20 | Day 100 | 3 | 51115 | 25481 |
| 6_PC_UN_OX_T0_3 | 16-06 | Unamended | 0-20 | Day 0 | 3 | 102921 | 47855 |
| 6_PC_UN_OX_T1_3 | 16-06 | Unamended | 0-20 | Day 50 | 3 | 58408 | 31579 |

